# The assembly of the membrane arm of mitochondrial complex I follows evolutionarily distinct routes

**DOI:** 10.1101/2025.06.10.658862

**Authors:** Simge Parlar, Michelle Marofke, Kristina Kühn, Ryan M.R. Gawryluk, Etienne H. Meyer

## Abstract

Complex I is the largest enzyme of the mitochondrial oxidative phosphorylation system. It is composed of core subunits that are conserved across bacteria and eukaryotes and accessory subunits that are lineage-specific. Consequently, the assembly of complex I likely requires lineage-specific assembly machineries. Complex I assembly has been mainly studied in mammals. It follows a modular pathway in which modules are assembled independently and then associated together to form the mature complex. Here, we identify and characterize PAF1 as an assembly factor for the membrane arm of complex I in plants. We show that this protein plays a role in stabilizing a module of the membrane arm prior to its association with the rest of the complex. PAF1 was likely present in the last eukaryotic common ancestor and has been lost in mammals in which the assembly of the membrane arm involves assembly factors not conserved in plants. Altogether, we provide molecular evidence that the assembly of the membrane arm of complex I can be performed by two different machineries.

## Introduction

Eukaryotes result from the endosymbiosis between an archaeon related to the Heimdallarchea (Eme et al, 2023) and a facultatively aerobic proteobacterium (Gray, 2012). They latter evolved into mitochondria, forming semi-autonomous organelles (Roger et al, 2017). During the evolution of eukaryotes, genes from the mitochondrial ancestor were mostly transferred to the nuclear genome or lost, and only a handful of genes remain on the mitochondrial genome (Adams & Palmer, 2003). The gene content of the mitochondrial genome varies between eukaryotic groups, highlighting the many evolutionary routes that were followed by mitochondrial lineages (Veeraragavan et al, 2024).

Mitochondria are double membrane-bound organelles. They are biochemical factories producing many key metabolites that are exported to the cytoplasm. These metabolites include several cofactors, building blocks such as organic and amino acids and energy in the form of ATP. ATP production is performed during a process called cellular respiration. In mitochondria, the TCA cycle is involved in the oxidation of substrates concomitantly with the production of reduced cofactors. These cofactors are recycled by the respiratory chain which transfers electrons via several protein complexes to molecular oxygen. During this electron transfer, a proton gradient is created across the mitochondrial inner membrane. The ATP synthase couples the diffusion of protons back into the matrix with ATP synthesis.

The main entry point of electrons into the respiratory chain is the NADH-ubiquinone oxidoreductase or complex I (Hirst, 2013, Parey et al, 2020). It oxidises NADH in the matrix and transfers electrons to ubiquinone. The pumping of protons into the inter membrane space is coupled with the electron transfer. Complex I is composed of 14 core subunits that are conserved across organisms. Additional, accessory subunits are present in many groups. Alphaproteobacterial complex I contains three accessory subunits (Yip et al, 2011). In eukaryotes, many additional subunits have been acquired; some are present in all eukaryotic lineages, whereas others are lineage-specific (Elurbe & Huynen, 2016, Padavannil et al, 2022). A few of the complex I subunits – all with counterparts in the bacterial complex I - are encoded by the mitochondrial genome, whereas the rest is nucleus-encoded. Complex I forms an L-shaped complex with one arm embedded in the inner membrane and the other arm protruding into the mitochondrial matrix (Hirst, 2013, Parey et al, 2020). Each arm is divided into two modules: the matrix arm is formed by the N-module and the Q-module, while the membrane arm is divided into the proximal (P_P_) and distal (P_D_) modules, based on their respective positions relative to the matrix arm (Hunte et al, 2010).

The assembly of complex I has been extensively studied in opisthokonts (i.e. animals, fungi, and their close unicellular relatives) in which detailed assembly models and mechanisms have been proposed based on genetic and structural studies (Laube et al, 2024, Sanchez-Caballero et al, 2016, Signes & Fernandez-Vizarra, 2018). Briefly, different assembly intermediates (some corresponding to the modules forming complex I) are first assembled separately and then they are associated together in a step-wise manner to form the mature complex (Guerrero-Castillo et al, 2017, Ligas et al, 2019, Meyer et al, 2019). Several assembly factors have been identified; they play various roles such as modification of a subunits or stabilisation of an assembly intermediates (Formosa et al, 2018, Laube et al, 2024, McKenzie & Ryan, 2010).

In plants, complex I is composed of 48 subunits, nine of which are encoded by the mitochondrial genome (Klusch et al, 2023). In comparison with bacterial or animal complex I, plant complex I contains two additional domains, a matrix-facing carbonic anhydrase domain anchored on the P_P_ module and a bridge domain linking the matrix arm and the carbonic anhydrase domain (Klusch et al, 2023, Klusch et al, 2021). The assembly of complex I in plants follows a modular pathway similar to the one in opisthokonts but the order of association of the assembly intermediates is different (Ligas et al, 2019, Meyer et al, 2022). In addition, many assembly factors identified in opisthokonts are not conserved in plants (Meyer et al, 2019) and additional assembly factors have been described in plants (Ivanova et al, 2019, Schertl et al, 2012, Schimmeyer et al, 2016). Altogether, these observations suggest that the assembly pathway of complex I in plants differs from the one described in opisthokonts.

In this study we identified a novel assembly factor in the model plant *Arabidopsis thaliana* that we named PAF1 for P_D_ module Assembly Factor 1. We propose that this protein is involved in the stabilization of the P_D_ module prior to its assembly. Sequence similarity searches suggest that PAF1 is not only present in plants but also in many other eukaryotic groups and not in animals. Overall, our results indicate that at least two different pathways exist for the assembly of the P_D_ module of complex I and we describe here an element, found in the Last Eukaryotic Common Ancestor (LECA), of the assembly machinery of the membrane arm of complex I.

## Results

### PAF1 is a mitochondrial protein present in most eukaryotic lineages

To identify proteins important for complex I proteostasis in *Arabidopsis thaliana*, we previously performed a bioinformatic search using gene function prediction tools (Hansen et al, 2018). One of the candidates identified is a small protein of 91 amino acids, encoded by the gene *At1g05205*, that we name here as PAF1 for P_D_ module Assembly Factor 1, containing no predicted transmembrane or functional domains. According to SUBA5 (Hooper et al, 2017), sub-cellular targeting predictions result in unclear outcomes but the protein encoded by *At1g05205* has been identified many times in proteomes of isolated mitochondria (Fuchs et al, 2020, Nietzel et al, 2020, Rugen et al, 2021, Senkler et al, 2017, Tan et al, 2012). Therefore, SUBA5 annotates PAF1 as a mitochondrial protein.

To gain more insights into the evolutionary history of PAF1, we searched for homologous proteins across the tree of life. PAF1-like proteins were found in many eukaryotic groups but not in prokaryotes (Figure 1A). Interestingly, eukaryotic groups that have lost mitochondria, such as metamonads, have also lost PAF1. PAF1 was also lost in most opisthokonts including all animals, and, among fungi, it is present in several groups, but not in Dikarya, the focus of nearly all fungal Complex I research. Based on this distribution, PAF1 appears to be an ancient eukaryotic protein, likely present in the LECA. Homologs of PAF1 are short proteins with poor sequence conservation (Figure 1B) except around the two cysteines predicted by Alphafold (Jumper et al, 2021) to form a disulphide bond at the base of a helix-loop-helix domain (Figure 1C).

**Figure 1.**
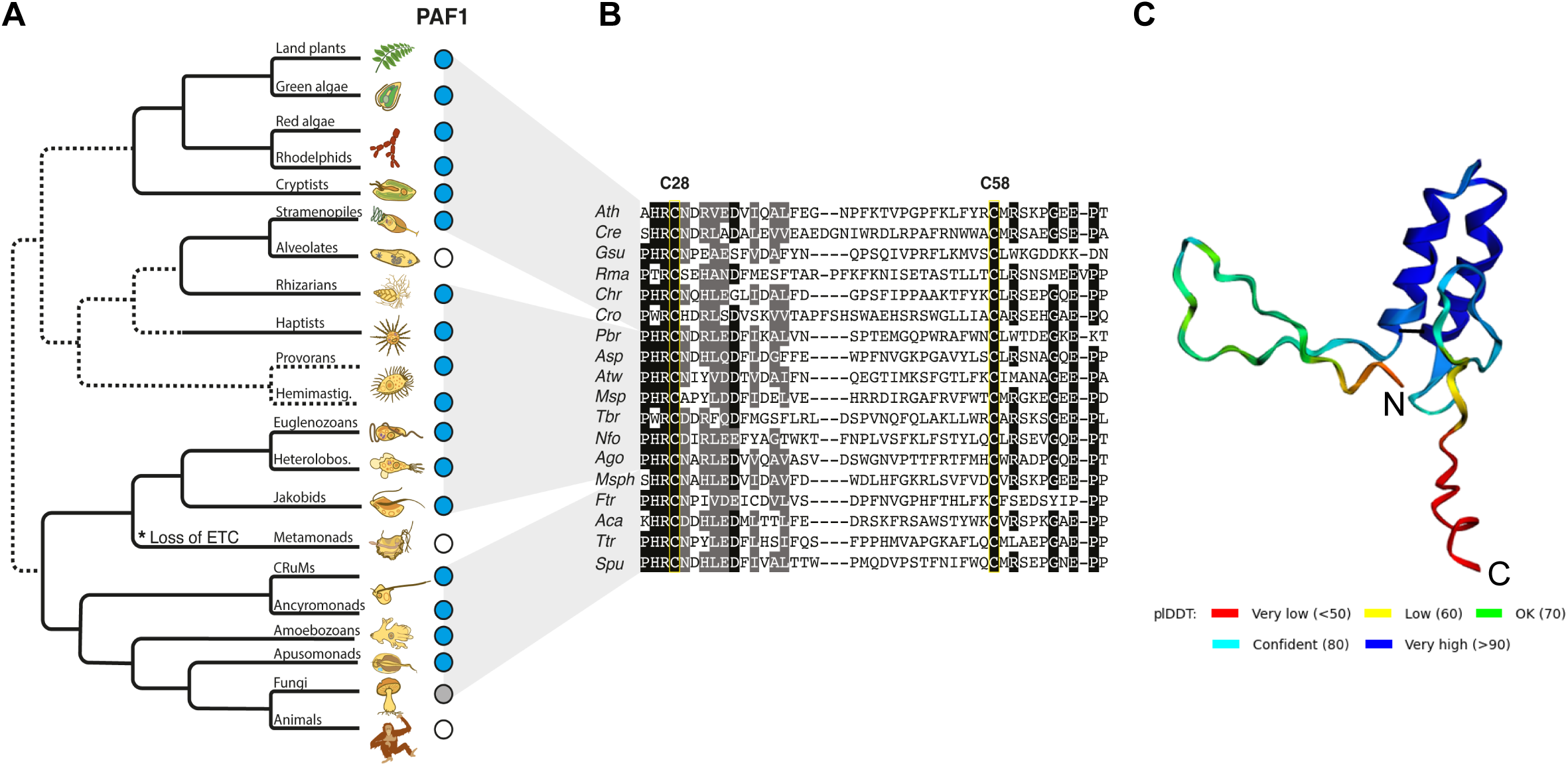
PAF1 is a small protein that was present in the last eukaryotic common ancestor. A. Phylogenetic tree schematic of eukaryotes demonstrating the widespread distribution and antiquity of PAF1. Unfilled and cyan circles indicate PAF1 absence and presence, respectively; however, presence does not necessarily mean that all or most lineages within a group encode PAF1. The grey circle associated with Fungi denotes that some groups encode PAF1, whereas dikaryons do not. Organismal drawings are from (Keeling & Eglit, 2023). The tree topology is a synthesis based on multiple phylogenomic analyses; dashed lines represent unresolved relationships. B. Phylogenetically broad sequence alignment of PAF1 homologs from diverse eukaryotes. A partial PAF1 sequence from a representative species of each eukaryotic group shown in A. is presented. The one-letter amino acid code is used, and residues highlighted in black indicate ≥ 70% identity at that site. Conserved disulphide bridge-forming cysteine residues corresponding to amino acids 28 and 58 in *Arabidopsis* PAF1 are highlighted by yellow boxes. Abbreviations: *Ath*, *Arabidopsis thaliana*; *Cre*, *Chlamydomonas reinhardtii*; *Gsu*, *Galdieria sulphuraria*; *Rma*, *Rhodelphis marina*; *Chr*, *Chroomonas b nsp.*; *Cro*, *Cafeteria roenbergensis*; *Pbr*, *Plasmodiophora brassicae*; *Asp*, *Acanthocystis* sp.; *Atw*, *Ancoracysta twista*; *Msp*, *Meteora sporadica*; *Tbr*, *Trypanosoma brucei*; *Nfo*, *Naegleria fowleri*; *Ago*, *Andalucia godoyi*; *Msph*, *Mantamonas sphyraenae*; *Ftr*, *Fabomonas tropica*; *Aca*, *Acanthamoeba castellanii*; *Ttr*, *Thecamonas trahens*; *Spu*, *Spizellomyces punctatus*. C. Structural model of PAF1 from *Arabidopsis* as given by Alphafold. The N and C-termini are indicated, the black line at the base of the helices indicates the predicted disulphide bridge. The model confidence is indicated by the colouring of the residues according to the pLDDT score.

### PAF1 is important for plant growth

To evaluate the function of PAF1, we obtained a T-DNA insertion mutant, containing a T-DNA inserted in the first intron of *At1g05205* (Figure 2A). Compared to the wild-type control Col-0, the *paf1* mutant shows a delayed growth phenotype during germination and early growth (Figure 2B) as well as at later growth stages (Figure 2C) but the mutant develops normally and slowly completes its life cycle. To confirm that the growth phenotype is caused by the absence of PAF1, we complemented the *paf1* mutant by reintroducing the native *PAF1* gene in the mutant background. The obtained lines are growing similarly to wild-type plants (Supp Figure 1). Altogether, this suggests that PAF1 is a small soluble mitochondrial protein important for plant growth.

**Figure 2.**
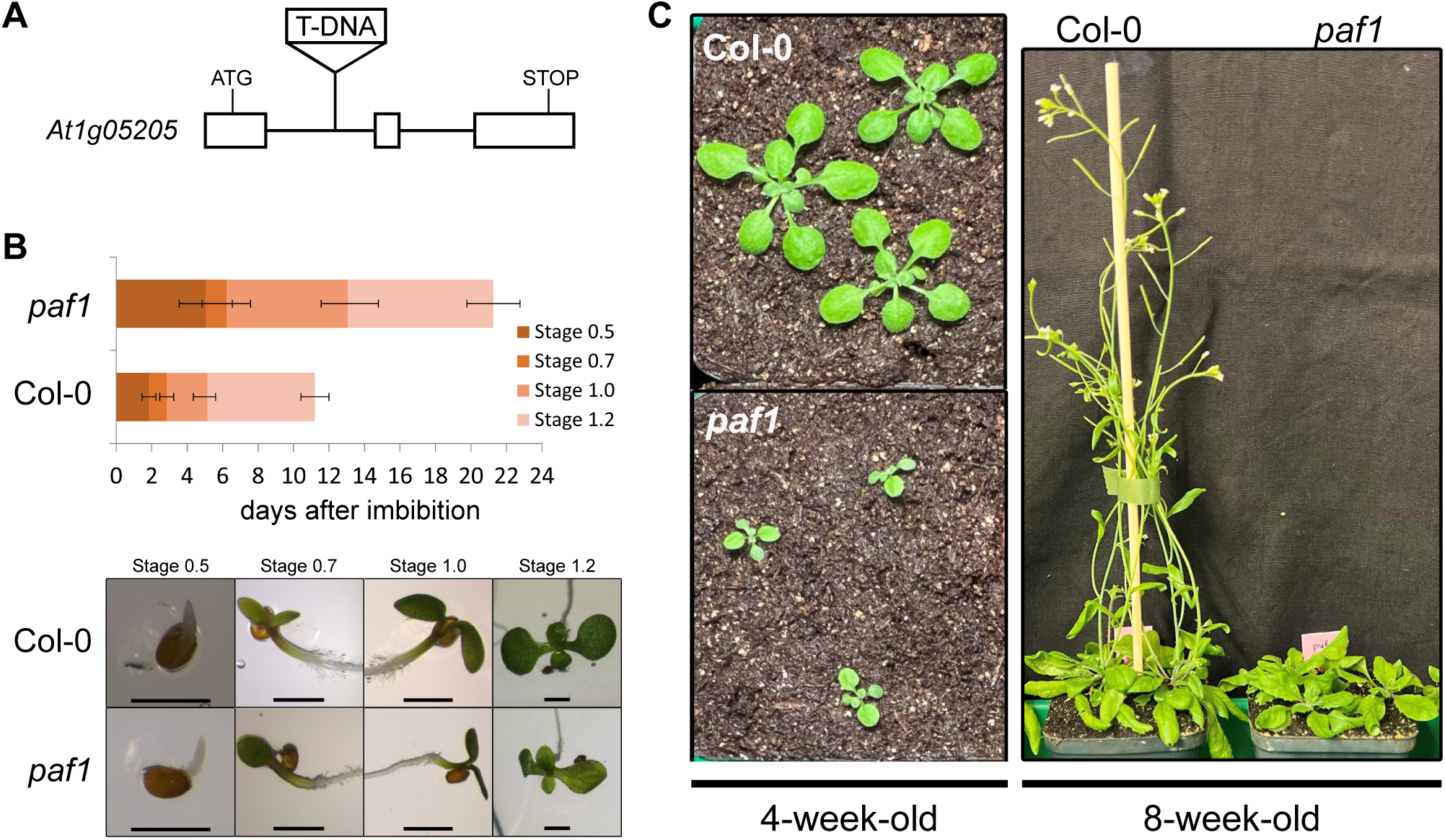
PAF1 is important for plant growth. A. Gene model of *At1G05205* encoding PAF1, the positions of the start and stop codons as well as the position of the T-DNA insertion are indicated. Boxes represents exons and lines introns. B. Analysis of early growth of *paf1*. The time needed to reach distinct developmental stages was recorded for *paf1* and Col-0 seedlings grown on synthetic medium, with stages according to (Boyes et al, 2001). Data are represented as mean plus standard deviation (n=81 for *paf1* and n=100 for Col-0). Lower panel, photo of a representative seedling at the corresponding stage. Scale bars represent 1 mm. C. Representative 4-week-old and 8-week-old plants of the different lines.

### PAF1 is important for complex I proteostasis

PAF1 was identified as a protein playing a putative role in complex I proteostasis (Hansen et al, 2018). To evaluate the influence of PAF1 absence on complex I levels, we extracted mitochondria from control and mutant lines and analysed protein complexes using native gel electrophoresis. In *paf1*, complex I is detected at much lower levels compared to the Col-0 control (Figure 3). In the two complementation lines, wild-type levels of complex I are restored (Figure 3), indicating that PAF1 is important for the maintenance of normal levels of complex I.

**Figure 3.**
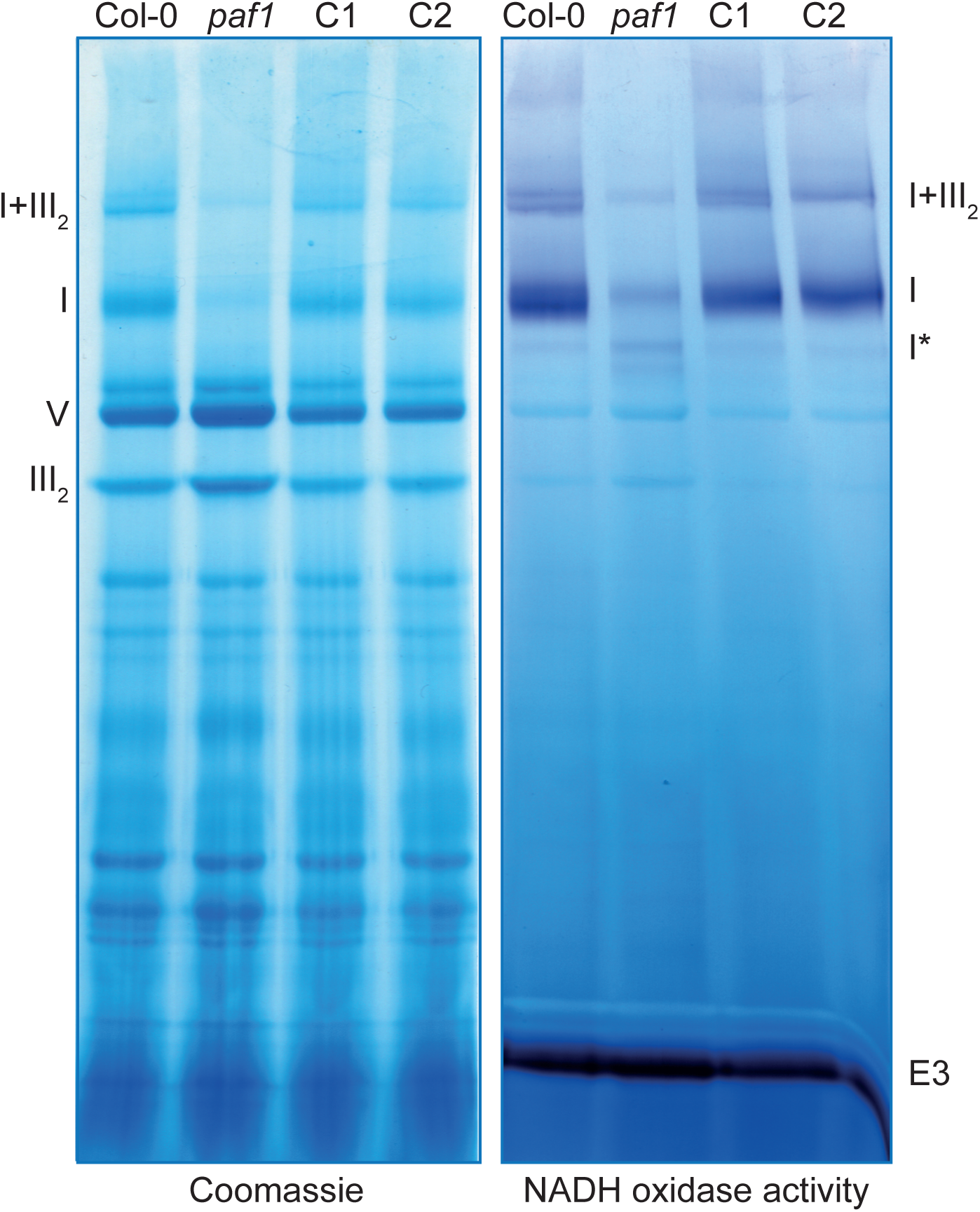
PAF1 is important for complex I proteostasis. Blue Native Polyacrylamide Gel Electrophoresis of mitochondrial complexes. Isolated mitochondria were solubilized with 5% digitonin and the solubilized complexes were separated in duplicate on a native gel. Half of the gel was stained with Coomassie blue (left panel) and the other half was stained for NADH oxidase activity (right panel). The localization of the OXPHOS complexes and supercomplexes is indicated on the sides (I: complex I, III_2_: complex III_2_, V: complex V, I+III_2_: supercomplex I+III_2_). E3: NADH oxidase activity of the E3 subunit of the pyruvate decarboxylase complex. C1 and C2 correspond to the two complemented lines.

To evaluate if the role of PAF1 is limited to complex I proteostasis, we performed an analysis of the total proteome of mitochondria isolated from *paf1*, Col-0 and *ndufv1*, a complex I mutant showing a developmental phenotype similar to *paf1* (Supp Figure 2). Statistical analysis of differences in protein abundances in the three samples show that the mitochondrial proteomes of *paf1* and *ndufv1* are almost identical whereas both mutants show proteome alterations when compared to Col-0 (Supp Figure 2). This suggests that the function of PAF1 is restricted to complex I proteostasis.

Another hint towards a complex I specific function of PAF1 comes from the analysis of *Viscum album*, a parasitic plant that lost complex I (Maclean et al, 2018, Senkler et al, 2018) and all the genes encoding complex I-specific proteins (Schroder et al, 2022). We were unable to find a homolog of PAF1 in the genome of *Viscum album*. The co-absence of PAF1 with complex I is consistent with a complex I-specific function.

To determine if PAF1 could be a previously unidentified complex I subunit, we mined published datasets. In the different structures of complex I from various plant species (Klusch et al, 2023, Klusch et al, 2021, Maldonado et al, 2023, Soufari et al, 2020), no additional densities potentially corresponding to PAF1 were observed. In complexome profiling data of mitochondrial complexes, PAF1 was identified in three datasets (Ligas et al, 2019, Rugen et al, 2021, Senkler et al, 2017) and was never identified in the fractions corresponding to complex I but rather in fractions corresponding to smaller complexes (Supp Figure 3). Taken together, these data exclude that PAF1 is a subunit of complex I and therefore it may act as a complex I assembly factor.

### PAF1 is important for the assembly of the P_D_ module

To gain further understanding on the role of PAF1, we evaluated the presence of complex I assembly intermediates in *paf1* using BN-SDS-PAGE. Antibodies directed against the subunit CA2 detected several complex I intermediates that overaccumulate in *paf1* (Figure 4A). CA2 is a subunit of the membrane that is assembled early (Ligas et al, 2019). Detection of CA2-containing intermediates suggests either impaired assembly of increased instability of complex I. The last stable assembly intermediate of complex I, complex I*, contains an assembly factor, L-galactono-1,4-lactone dehydrogenase (GLDH). The detection of GLDH can be used to discriminate between assembly and stability defects as the main breakdown product of complex I is an intermediate similar to complex I* but lacking GLDH (Maldonado et al, 2020). Using anti-GLDH antibodies, we detected the accumulation of complex I* (Figure 4B), indicating that the assembly of complex I is impaired in *paf1*. GLDH is bound to complex I* (Schertl et al, 2012, Schimmeyer et al, 2016, Soufari et al, 2020) and needs to be released before the distal module of the membrane arm (P_D_ module) can be assembled (Ligas et al, 2019, Meyer et al, 2022). In comparison to Col-0, lower amounts of free GLDH are detected in *paf1* (Figure 4B). Altogether, this suggests that the release of GLDH and the association of the P_D_ module on complex I* are impaired in the absence of PAF1. Such an assembly defect could lead to the accumulation of the unassembled P_D_ module. To test this hypothesis, we used antibodies directed against Nad5 and NDUB9, two subunits of the P_D_ module. In Col-0, Nad5 is detected at the level of complex I and supercomplex I+III but also in a low abundance complex of ca. 400 kDa, located in between two signals obtained with the anti-CA2 antibodies (Figure 4C). In *paf1*, no accumulation of Nad5-containing intermediates is detected (Figure 4C). In both lines, NDUFB9 is only detected in complex I and the supercomplex I+III (Figure 4D), indicating that this subunit is assembled very late as previously suggested (Ligas et al, 2019). This analysis demonstrates that, in the absence of PAF1, the assembly of complex I, in particular the formation and the association of the P_D_ module to complex I*, is impaired.

**Figure 4.**
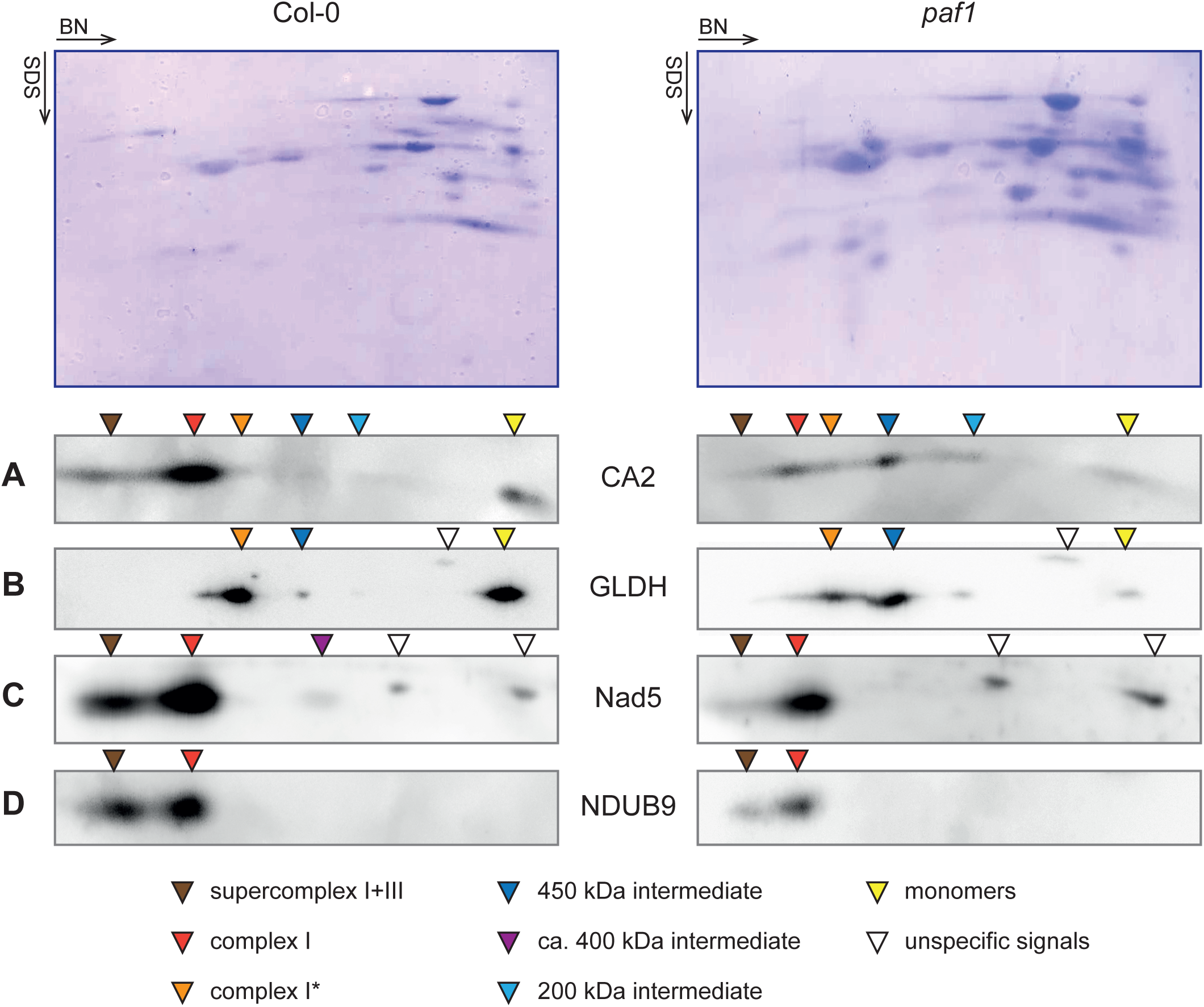
PAF1 is important for the assembly of the P_D_ module. Isolated mitochondria were solubilized with 5% digitonin and the solubilized complexes were separated on a native gel. The subunits were further resolved in a second dimension using SDS-PAGE and transferred on membranes that were probed with several antibodies. Top panels show representative images of the membranes stained with Coomassie blue. A: Signals obtained using the anti-CA2 antibodies. B: Signals obtained using the anti-GLDH antibodies. C: Signals obtained using the anti-Nad5 antibodies. D: Signals obtained using the anti-NDUB9 antibodies. The position of the detected complexes is indicated with coloured triangles.

### PAF1 is not involved in mitochondrial gene expression

As a soluble mitochondrial protein, it is plausible that PAF1 may be involved in the expression of the two mitochondria-encoded subunits of the P_D_ module, Nad4 and Nad5. Impaired expression of Nad4 and Nad5 lead to the accumulation of the 450 kDa intermediate and complex I* (Ligas et al, 2019), assembly defects similar to those observed in *paf1* (Figure 4). To evaluate mitochondrial gene expression in *paf1*, we extracted total RNA and performed RNA gel blot hybridizations to detect the levels of selected mitochondrial transcripts. We did not detect any alteration of transcript patterns between *paf1* and Col-0 (Figure 5). This indicates that PAF1 is not involved in mitochondrial gene expression.

**Figure 5.**
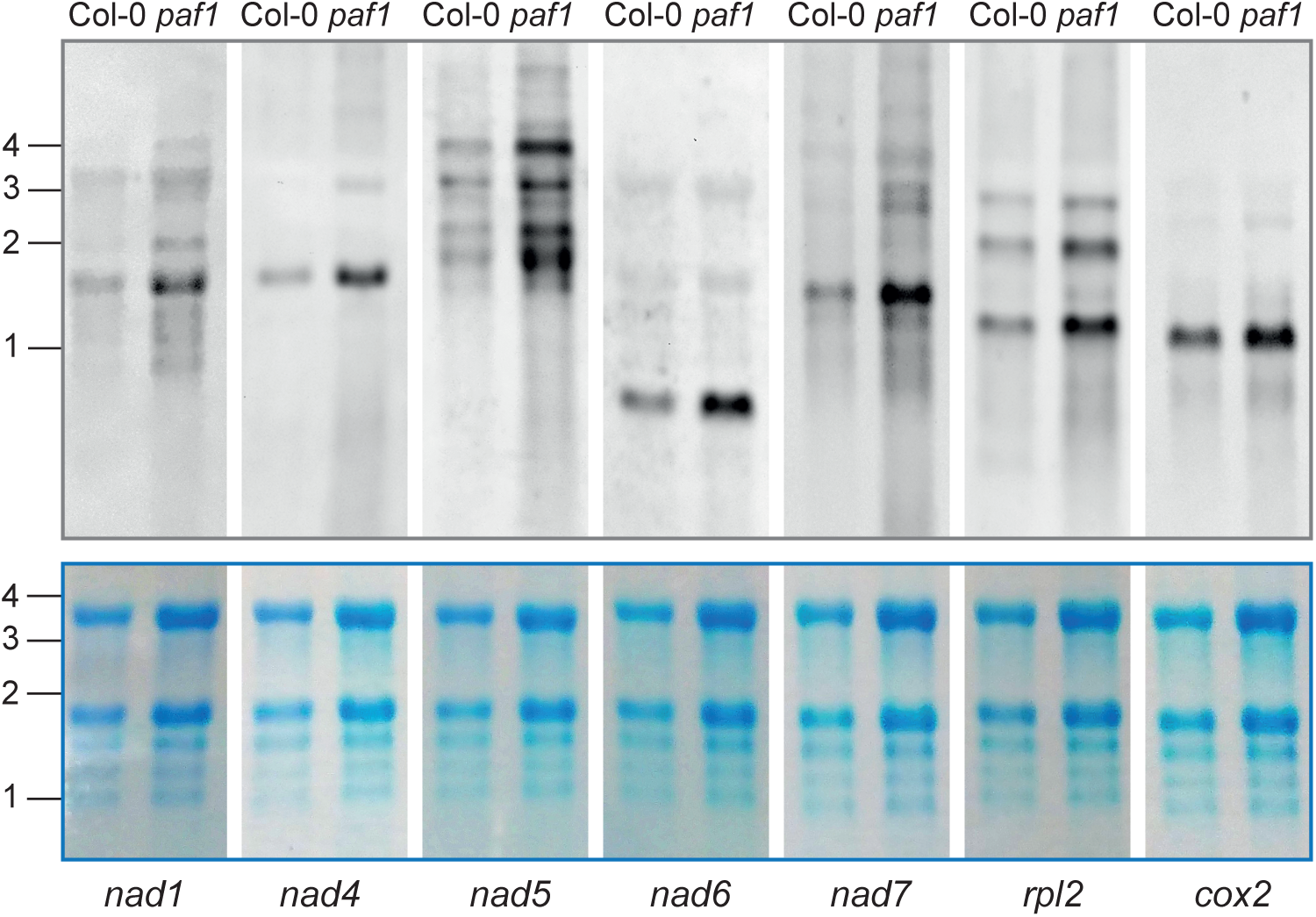
PAF1 is not involved in mitochondrial gene expression. Total RNAs were extracted and separated on a denaturing agarose gel. The RNAs were transferred on membranes that were incubated with probes specific for selected mitochondrial transcripts. Top panels: Signals obtained using the different probes. Bottom panels: Methylene blue staining of the corresponding membranes. Sizes in kb are indicated on the left of the panels. The transcripts probed are indicated below the panels.

### PAF1 interacts with Nad4

Antibodies against Nad5 detected a complex at ca. 400 kDa in Col-0 but not in *paf1* (Figure 4C). To gain further insights into this complex and evaluate if it corresponds to the P_D_ module of the membrane arm of complex I, we analysed the data corresponding to the lower part of BN gels in complexome profiling datasets. In the dataset from Rugen et al (Rugen et al, 2021), the subunits of the P_D_ module cluster together in a complex of ca. 400 kDa (Supp Figure 4). In the dataset from Senkler et al (Senkler et al, 2017), fewer complex I subunits are detected in the lower part of the gel but several of the subunits of the P_D_ module are also found in a complex of ca. 400 kDa (Supp Figure 5). Therefore, the P_D_ module is detectable in Col-0 with the anti-Nad5 antibodies but it does not accumulate in *paf1*. In the dataset from Rugen et al, PAF1 is detected in the same area of the gel, suggesting that it is associated with the P_D_ module (Supp Figure 4). In the dataset from Senkler, PAF1 is not detected in the P_D_ module but it is found together with Nad4 in a lower molecular weight fraction (Supp figure 5). This analysis suggests that PAF1 is interacting with the P_D_ module, most likely at the level of subunit Nad4.

To gain further insights into the putative interaction of PAF1 with subunits of the P_D_ module, we performed structural predictions using Alphafold2 (Jumper et al, 2021). An interaction between PAF1 and Nad4 is predicted with high probability (Figure 6A) whereas interactions with Nad5 or Nad2, the subunit of the P_P_ module that interacts with Nad4, are not predicted (Supp Figure 6). The confidence of the prediction of the N-terminal domain of PAF1 increases in the presence of Nad4 (plDDT > 80, Figure 6A) as compared to when PAF1 folding is predicted alone (plDDT = 70, Figure 1C), suggesting that the N-terminal domain of PAF1 interacts with Nad4. PAF1 is predicted to bind on the intermembrane-space side of Nad4, with its N-terminal domain at the position where the subunit NDUP1 binds in the fully assembled complex I (Figure 6B). As PAF1 is conserved in several eukaryotic lineages, its role in complex I assembly must also be conserved. Therefore, PAF1 homologs in the different eukaryotic lineages should interact with the corresponding Nad4 homolog. We performed structural predictions to evaluate the interactions between PAF1 and Nad4 in six eukaryotic groups. When a confident interaction is predicted (four out of six cases), PAF1 is predicted to interact with Nad4 on its IMS side, at the same position as predicted in Arabidopsis (Supp Figure 7). Therefore, based on proteomic and structural modelling data, we conclude that PAF1 binds on the IMS side of Nad4.

**Figure 6.**
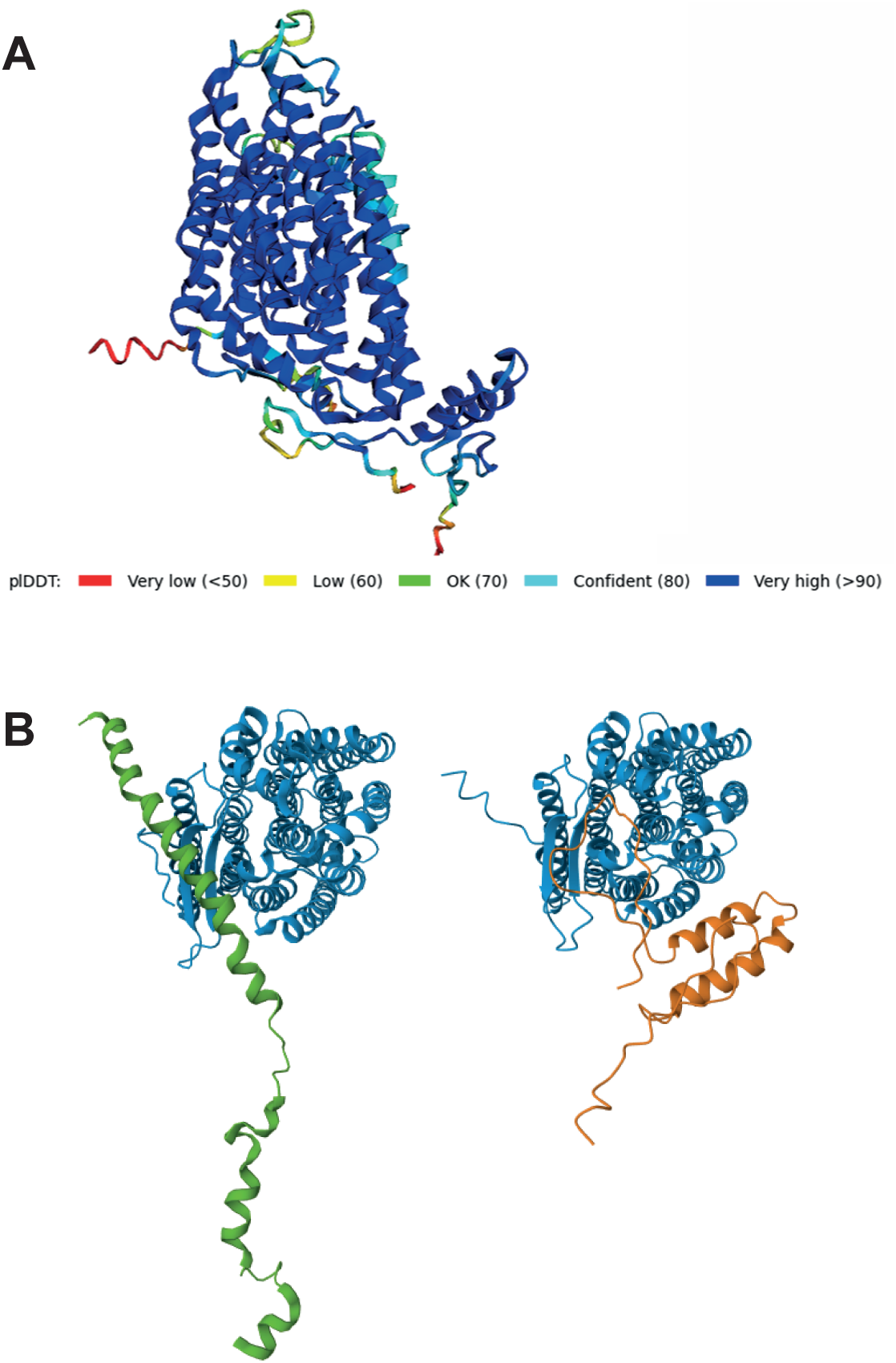
PAF1 is predicted to interacts with Nad4. A: Prediction of the interaction between PAF1 and Nad4 using Alphafold2 (multimer option). The full-length sequences of the proteins were utilized and default parameters were used. The plDDT score is given by Alphafold to indicate the confidence of the model and is shown using a colour scale. The view is from the membrane with the matrix side on top on the IMS side at the bottom of the image. B. Comparison of the interaction between Nad4 and NDUP1 in the complex I structure and the interaction model between PAF1 and Nad4. The structure of Nad4 and NDUP1 were extracted from the model of complex I published by Klusch et al. 2023 (PDB: 8BPX). Both structure and model are shown as a view from the IMS.

## Discussion

In this study, we identified and characterized PAF1, an assembly factor important for the assembly of the membrane arm of complex I. Our data indicate that PAF1 binds to subunit Nad4 of the P_D_ module of the membrane arm. In the absence of PAF1, the P_D_ module of complex I is not detectable and complex I*, the assembly intermediate to which the P_D_ module attaches, accumulates. Altogether our data suggest that PAF1 stabilizes the P_D_ module prior to its association with complex I*.

The assembly of the P_D_ module has been proposed to act as a check-point for the assembly of complex I in plants (Meyer et al, 2022). This step appears to be the limiting step of the assembly pathway as the last assembly intermediate, complex I*, containing the matrix arm attached to the P_P_ module, is detected to low levels under normal conditions (Ligas et al, 2019). Complex I* contains an assembly factor, GLDH, that is bound on the IMS side of subunit Nad2 at the position where the subunit NDUP1 is found in the mature complex (Meyer et al, 2022). GLDH must then be detached to allow the assembly of subunit NDUP1, a subunit that is spanning the membrane arm on the IMS side similar to a ‘piece of tape’ locking the P_P_ and P_D_ modules together (Klusch et al, 2023, Maldonado et al, 2023, Soufari et al, 2020). In *paf1*, GLDH is found mostly associated with complex I assembly intermediates, suggesting that it cannot be detached from complex I* and its precursors. As we observed that the P_D_ module does not accumulate in *paf1* (Figure 4), our data suggest that it is the association of the P_D_ module that releases GLDH from Complex I* rather than GLDH being detached prior to the association of the P_D_ module. PAF1 is predicted to bind to the IMS side of Nad4 at the position where NDUP1 is located in the mature complex I (Figure 6). Therefore, similar to GLDH, PAF1 must also be released from the P_D_ module upon association with complex I* to allow the binding of NDUP1 on the assembled membrane arm. Consequently, the last step of complex I assembly in plants involves the association of the PAF1-containing P_D_ module with complex I*, an event linked with the release of both GLDH and PAF1 to free the binding sites for NDUP1 both on Nad2 (P_P_ module) and Nad4 (P_D_ module).

We observed that the P_D_ module migrates at an apparent size of ca. 400 kDa (Figure 4). This size is similar to the one observed in complexome profiling experiments (Supp Figures 4 and 5). However, the calculated molecular weight of the P_D_ module is much smaller (ca. 230 kDa). This suggests that the P_D_ module likely dimerizes before its assembly. We did not detect any accumulation of Nad5-containing intermediates such as a putative P_D_ module monomer in *paf1*. This suggests that either PAF1 is not involved in the dimerization or that the monomer is highly unstable. A similar apparent size compatible with the dimerization of the P_D_ module has been observed in animals (Guerrero-Castillo et al, 2017). This suggests that the dimerization of the P_D_ module prior to its assembly is an intrinsic property of this domain that is conserved across species.

We found that PAF1 is a mitochondrial protein that was present in LECA. Interestingly it has been lost in most opisthokonts, the best studied organisms for complex I assembly. In mammals, several assembly factors have been identified as important for the assembly of the membrane arm. Most of these are not conserved in plants (Meyer et al, 2019). Similarly, the two assembly factors characterized in plants for the last step of the assembly of the membrane arm of complex I, GLDH and PAF1, are not conserved in mammals. In mammals, four assembly factors have been identified as important for the assembly of the distal part of the membrane arm. TMEM70 is an assembly factor of the mitochondrial ATP synthase that also interacts with and stabilizes assembly intermediates of complex I, in particular the P_D_ module (Sanchez-Caballero et al, 2020). A protein homologous to TMEM70 is encoded by the Arabidopsis genome but its function has not been characterized yet. DMAC1 and ATP5SL/DMAC2 have been shown to interact with newly translated ND5 and several subunits of the P_D_ module (Stroud et al, 2016). In addition, in the absence of these assembly factors, a late-stage assembly defect is observed and therefore DMAC1 and DMAC2 were proposed to be important for the assembly of the P_D_ module (Stroud et al, 2016). FOXRED1 has been identified as an assembly factor involved in a late-stage assembly event (Formosa et al, 2015). Later, FOXRED1 was found enriched with DMAC1 and DMAC2, suggesting that they are all involved in the same assembly step (Stroud et al, 2016). These three proteins are not conserved in plants. DMAC1 appears to be a vertebrate invention whereas FOXRED1 homologs are founds in various prokaryotic and eukaryotic lineages (Lemire, 2015).

The pathway we propose involves the assembly factors PAF1 and GLDH. Apart from its role in complex I assembly, GLDH is an enzyme involved in ascorbic acid (vitamin C) biosynthesis in the pathway described for photosynthetic organisms (Smirnoff & Wheeler, 2000, Wheeler et al, 2015). However, homologs of GLDH are found in many non-photosynthetic organisms (Wheeler et al, 2015), indicating that GLDH was also likely present in LECA. Therefore, two distinct mechanisms have evolved for the assembly of the membrane arm of complex I: an ancestral pathway involving GLDH and PAF1 and a more recent pathway that evolved in opisthokonts. Further work is required to better characterize these two assembly pathways.

The discovery that the assembly of the membrane arm of complex I involves ancestral proteins confirms that plant mitochondria followed a different evolutionary route than animal mitochondria did. In particular, plant mitochondria utilize ancestral features for the assembly of their OXPHOS complexes whereas animal mitochondria have replaced these with relatively simpler pathways. The maturation of *c*-type cytochromes involves many proteins in plants but only two in animals (Allen, 2011). The transfer of the Rieske-Iron-Sulphur protein into complex III is performed by an ancestral Twin-Arginine Transport pathway in plants (Schäfer et al, 2020) and by a single BCS1 protein in animals (Wagener & Neupert, 2012). The formation of the membrane arm of complex I involves several ancestral subunits (e.g. carbonic anhydrases) and assembly factors (GLDH, PAF1) that have also been lost in animals where a new set of assembly factors have been recruited. However, it remains to be evaluated if the assembly of the membrane arm of complex I in animal is simpler than in plants as both pathways are not fully characterized yet. Overall, the fact that less proteins are involved in these assembly steps in animals suggests that animal mitochondria have evolved to optimize the cost of biogenesis. Such an evolutionary path has not been followed by plant mitochondria, most likely because plants are producing their own food via photosynthesis and therefore plant mitochondria rarely encounter substrate limitation.

In conclusion, we identified and characterized an assembly factor of the membrane arm of complex I in plants. This assembly factor is an ancient eukaryotic invention that was likely an assembly factor of complex I in LECA. In animals, it has been lost, most likely together with the ancestral machinery for the assembly of the membrane arm of complex I. This indicates that several mitochondrial pathways evolved to perform the same task and therefore characterizing mitochondrial biogenesis in various eukaryotic lineages might identify additional machineries.

## Methods

### Sequence alignments and phylogenetic analysis

An iterative multi-step search process was used to identify putative homologs of PAF1 in diverse eukaryotes. Initially, obvious homologs of PAF1, primarily from non-flowering plants, green algae, and several other groups were identified by iterative PSI-Blast searches at NCBI. These homologs were aligned with MAFFT, and a profile Hidden Markov Model (HMM) was generated with HMMER3. HMM searches were used to interrogate databases containing predicted protein sequences from diverse eukaryotes, with an emphasis on microbial eukaryotes. Putative homologs were screened manually. Briefly, those identified typically had e values less than 10^-3^ and were similar in size and physicochemical characteristics to Paf1. When new high confidence homologs were identified, the HMM was updated to include them, and further searches were performed as described above.

### Establishment of plant lines

The T-DNA lines SALK_ 020381 was obtained from the Nottingham Arabidopsis Stock Center and homozygous seedlings were obtained after selfing and confirmed by genotyping. For the complementation, the genomic region corresponding to the promoter and the coding sequence of PAF1 was amplified by PCR using primers paf1_pro-F and paf1_stopR and cloned between the KasI and EcoRI sites of the pORE_E3 vector (Coutu et al, 2007) that had been modified by replacing the *Basta^R^* gene by the *Hygromycin^R^* gene. The resulting vector was transformed into *Agrobacterium tumefaciens* GV3101 and this strain was used to transform the T-DNA line by floral dip. Transformants were screened on hygromycin. A hygromycin sensitivity test was performed on T3 plants to identify complemented lines homozygous for the complementation construct. The sequences of all the primers used are given in Supp Table S1.

### Plant cultivation

Seeds were surface sterilized with ethanol 70% (v/v) containing 0.05% (v/v) tween20 and plated on MS media containing 1% (w/v) sucrose. Plates were incubated under long day conditions (16h light 120 µE, 22°C and 8h darkness, 20°C, the humidity was not controlled and oscillated between 40 and 65%). 10-day-old seedlings were transferred on soil and grown under the same long day conditions. The growth stage analysis was performed while the seedlings were on the plates.

### Mitochondria isolation

Mitochondria were isolated from 4-week-old plants according to (Meyer et al, 2009). Briefly, the aerial parts of the plant were ground in extraction buffer (0.3 M sucrose, 15 mM potassium pyrophosphate, 2 mM EDTA, 10 mM KH_2_PO_4_, 1% (w/v) polyvinylpyrollidone-40, 1% (w/v) bovine serum albumin, 20 mM sodium ascorbate, pH 7.5) in a cold mortar. After filtration, two rounds of differential centrifugations (2.000 g/20.000 g) were performed to obtain a mitochondria-enriched pellet. After resuspension in wash buffer (0.3 M sucrose, 1 mM EGTA, 10 mM MOPS/KOH pH 7.2), the fraction was loaded on a discontinuous Percoll gradient comprised of three layers (18%-25%-50%) which was centrifuged at 40.000 g for 45 min. The mitochondria were collected at the interface between the 25% and 50% layers and the Percoll was eliminated by performing several washes in wash buffer. Protein concentration was estimated using the Bradford assay (Rotiquant, Roth).

### Gel electrophoresis and western blot

For the BN-PAGE, 100 µg of mitochondrial proteins were treated with 5% (w/v) digitonin and the solubilized protein complexes were separated on a BN gel according to (Eubel et al, 2005). After migration, the complexes were stained using colloidal coomasie. For the BN-SDS-PAGE, gel lanes from the BN gel were excised and incubated in 1% (w/v) SDS and 1% (v/v) βmercaptoethanol for 30 min at RT. They were then placed on top of a 12% SDS-poly acrylamide gel (Laemmli, 1970). After the migration, the proteins were transferred on PVDF membrane (Immobilon-P, Millipore) using a wet blotting system. After transfer, the membranes were stained for 5 min in staining buffer (30% (v/v) methanol, 7% (v/v) acetic acid, 0.005% (w/v) Coomassie Blue R250). The background was reduced by washing the membranes in 40% (v/v) methanol, 10% (v/v) acetic acid. The stained membranes were scanned and the staining was fully removed using 100 % (v/v) methanol.

### Antibody production and western blot

Polyclonal antibodies were produced in rabbits against the peptides HAMSDEQDMRKMGGLASS and EVSFEALDKGAIEILGP specific to Nad5 (Biotem, France). The final bleed was used for western blots experiments. Membranes were first blocked using a 5% milk solution and incubated with the primary antibodies for 16h at 4°C. Following three washes with TBS-T (20 mM Tris-HCl pH 7.4, 150 mM NaCl, 0.1 % (v/v) Tween 20), the membranes were incubated with the secondary antibodies conjugated with HRP for 1h at 4°C and again washed three times with TBS-T. Final detection was performed using the ECL prime detection reagent (Cytiva Amersham) and a CCD camera (Fusion FX, Vilber). The primary antibodies were used at the following dilutions: anti-CA2 (Rohricht et al, 2023) 1/10000, anti-GLDH (Agrisera AS06182) 1/5000, anti-Nad5 1/5000 and anti-NDUB9 (PhytoAB PHY1085A) 1/5000.

### RNA Gel Blot Analysis

RNA extraction was performed on 7-10 days old Arabidopsis seedlings using TRIzol™ Reagent (Invitrogen) according to the manufacturer’s protocol. RNAs were additionally purified using Phenol-Chloroform-Isoamyl alcohol 25:24:1 (Roth) followed by an ethanol precipitation. 2 µg of total RNAs were resolved on 1.2 % (w/v) agarose/formaldehyde gels. RNA gel blots were performed as described by (Gualberto et al, 2015). DNA probes were generated using Arabidopsis genomic DNA or cDNA as template, the PCR DIG probe synthesis kit (Roche Applied Science) and primer pairs listed in Supp Table S1. RNA probes were transcribed from PCR products using T7-RNA-Polymerase (Thermo Fisher Scientific) and DIG RNA labeling mix (Roche Applied Science).

### Proteomic analysis

#### Protein digestion

Proteins from isolated mitochondria (10 µg of protein per sample) were dissolved in solubilization buffer (6 M urea/2 M thiourea pH 8.0) to adjust the protein concentration to 0.5 µg/µl. Proteins were reduced by incubation for 30 min at 22°C with DTT (1 µg per 10 µg protein) and then alkylated by incubation for 20 min at 22°C in the dark with iodoacetamide (5 µg per 10 µg protein). Samples were diluted four times with 10 mM Tris-HCl pH 8.0 and the proteins were digested for 15 h at 37°C using a Trypsin/LysC mixture (Promega, 0.4 µg protease per 10 µg protein). The samples were acidified to 0.2 % (v/v) TFA and centrifuged at 14 000 g for 10 minutes to eliminate any insoluble material. The peptides present in the supernatant were purified using ZipTips with C18 resin (Millipore) according to the manufacturer instructions and dried down.

#### MS analysis

The peptides were resuspended in 3 % (v/v) acetonitrile, 0.1 % (v/v) formic acid and 150 ng of digested peptides were separated on a Dionex UltiMate 3000 RSLCnano system consisting of a NCS-3500RS Nano ProFlow and WPS-3000TPL RS UPLC System (ThermoFischer, Dreieich, Germany) equipped with a PepMap™ Neo 5 µm C18 300 µm × 5 mm Trap cassette (ThermoFischer, Dreieich, Germany, https://www.thermofisher.com/order/catalog/product/de/de/174500) as well as a ACQUITY UPLC M-Class Peptide BEH C18 Column, 130Å, 1.7 µm, 75 µm × 250 mm (Waters, Eschborn, Germany, https://www.waters.com/nextgen/de/de/shop/columns/186007484-acquity-uplc-m-class-peptide-beh-c18-column-130a-17--m-75--m-x-2.html). After peptides were trapped for 10 min at 15 μl/min at 3% solvent B (solvent A: 0.1% (v/v) formic acid in water, solvent B: 0.1% (v/v) formic acid in ACN) and separated at 240 nl/min with a linear gradient of 3-50% B within 50 min. Eluting peptides were ionized at 2.1 kV and 320 °C in a Exploris 480 ThermoFischer nanoESI source using a Pre-cut PicoTip Emitter; 360 μm OD × 20 μm ID, 10 μm tip; 12 cm long (CoAnn Technologies, LLC). The Exploris 480 was equipped with a FAIMS device. The settings used were: CV of-50 V, MS range of 350-1500 m/z, 1 s cycle time, 120000 Orbitrap resolution, with a normalized AGC target of 300%, and minimum IT of 50 ms. Centroided MSMS of each selected precursor were collected at normalized HCD of 31% and a resolution of 15000, an AGC target of 100% and IT of 35 ms. Each precursor was then excluded from analysis for 30 s after one hit.

#### Data analysis

Acquired LC-MS/MS spectra were queried against an in-house modified Araport11 database including all nuclear encoded proteins as well as protein models of the mitochondrial and plastid genomes after RNA editing using MaxQuant 2.1.3.0 (Cox & Mann, 2008). Default parameters were used, and the protein quantification function was selected. The output proteingroup file was then imported into Perseus 1.6.15.0 (Tyanova et al, 2016) and the LFQ values were used for calculations. Differentially abundant proteins (DAP) between two genotypes were identified by performing Student’s t-test (two-tailed; FDR: 0.05, S0: 0.1) and visualized using the volcano plot option of Perseus.

The mass spectrometry proteomics data have been deposited to the ProteomeXchange Consortium via the PRIDE (Perez-Riverol et al, 2025) partner repository with the dataset identifier PXD064629.

### Structural predictions

Structural predictions were performed using ColabFold v1.5.5 (Mirdita et al, 2022) using default parameters.

## Acknowledgments

We thank Michael Kunzler and Felix Meyer for their technical help in isolating and characterizing the complementation lines. The work was financially supported by a grant of the Deutsche Forschungsgemeinschaft (ME 4174/3-1) to EHM. RMRG was supported by a grant from the Natural Sciences and Engineering Research Council of Canada (RGPIN-2019-04336).

## Author Contributions

Project design: EHM; Experimental work: SP, MM, MK, RG, EHM; Data analysis: SP, MM, KK, RG, EHM; Paper writing: EHM with contributions from all authors.

## Disclosure and competing interests statement

The authors declare no competing interests.

## Data availability

Proteomics data are available via ProteomeXchange with identifier PXD064629.

## Reviewer access details

Username

Password: 4ic5klmGJgB5

## Supplemental information

**Supp Figure S1: Complementation of *paf1* restored plant growth**

A. Schematic representation of the T-DNA used to complement the paf1 mutant.
B. Representative 6-week-old plants of the different lines. Two independent complementation lines were kept for analysis and named C1 and C2.
C. Analysis of early growth of the complemented lines. The time needed to reach several developmental stages was recorded for each line. Data are represented as mean plus standard deviation (n=21 to 36).

**Supp Figure S2: Analysis of the mitochondrial proteomes of *paf1* and *ndufv1***

A: Comparison of the growth phenotype of the ndufv1 and paf1 mutants. Top image: 5-week-old plants

Bottom image: 9-week-old plants.

B-D: Volcano plot representing the variation of the mitochondrial proteomes of Col-0, paf1 and ndufv1. Mitochondria were isolated from the three lines in triplicates and the mitochondrial proteomes analysed using mass spectrometry. Label-free quantification was performed and the difference in abundance of mitochondrial proteins were calculated using Perseus. Each square represents a protein. Filled circles present above the black line indicate proteins whose abundance is significantly changed between the two genotypes compared (t-test, FDR:0.5, S0:0.1). B. Comparison of the proteomes of paf1 and ndufv1. C. Comparison of the proteomes of paf1 and Col-0. D. Comparison of the proteomes of ndufv1 and Col-0.

**Supp Figure S3: Complexome profiling analysis of the association of PAF1 with mitochondrial complexes**

Complexome profiling datasets were mined for the presence of PAF1. Data were extracted from published datasets either via www.complexomemap.de or as previously published own work. The complex I subunits (CA2, NDUS1, Nad7 and NDUA3) were included to serve as positive control and show the position of complex I at 1000 kDa. A. Dataset from Rugen et al. B. Dataset from Senkler et al. C. Dataset from Ligas et al.

**Supp Figure S4: Analysis of the complexome profiling experiment published by Rugen et al.**

The leaf complexome profiling dataset was extracted from www.complexomemap.de. To visualize better the low abundant intermediates, the fractions corresponding to gel portions above 500 kDa were excluded. The molecular weight markers (in kDa) from www.complexomemap.de are indicated on the top. For each protein, the abundance in each fraction is represented using a color code: dark blue indicates the fraction in which the protein is the most abundant (in this part of the gel) and white indicates that the protein was not identified in the respective fraction. A hierarchical clustering was applied using Nova (default parameters). All the complex I subunits detected are included as well as PAF1. The cluster tree is indicated on the left. Subunits names are highlighted according to the module they belong to: Yellow: Q module, Pink: N module, Dark Green: Bridge domain, Blue: P_P_ module including the CA domain (light blue) and red: P_D_ module. PAF1 is highlighted in green.

**Supp Figure S5: Analysis of the complexome profiling experiment published by Senkler et al.**

For each protein, the abundance in each fraction is represented using a color code: dark blue indicates the fraction in which the protein is the most abundant (in this part of the gel) and white indicates that the protein was not identified in the respective fraction. A hierarchical clustering was applied using Nova (default parameters). All the complex I subunits detected are included as well as PAF1. The cluster tree is indicated on the left. Subunits names are highlighted according to the module they belong to: Yellow: Q module, Pink: N module, Dark Green: Bridge domain, Blue: P_P_ module including the CA domain (light blue) and red: P_D_ module. PAF1 is highlighted in green.

**Supp Figure S6: Structural predictions of potential interactions between PAF1 and Nad5 or Nad2**

Predictions were performed with Alphafold 2 using the multimer option. The full-length sequences of the proteins were utilized and default parameters were used. A: Prediction of the interaction between PAF1 and Nad5. B: Prediction of the interaction between PAF1 and Nad2. pLDDT: model confidence given by Alphafold.

**Supp Figure S7: Structural predictions of potential interactions between PAF1 and Nad4 in different eukaryotic lineages**

Predictions were performed with Alphafold 2 using the multimer option. The full-length sequences of the proteins were utilized and default parameters were used. The interaction between PAF1 and Nad4 in Arabidopsis is shown as reference. pLDDT: model confidence given by Alphafold.

**Supp Table S1:**
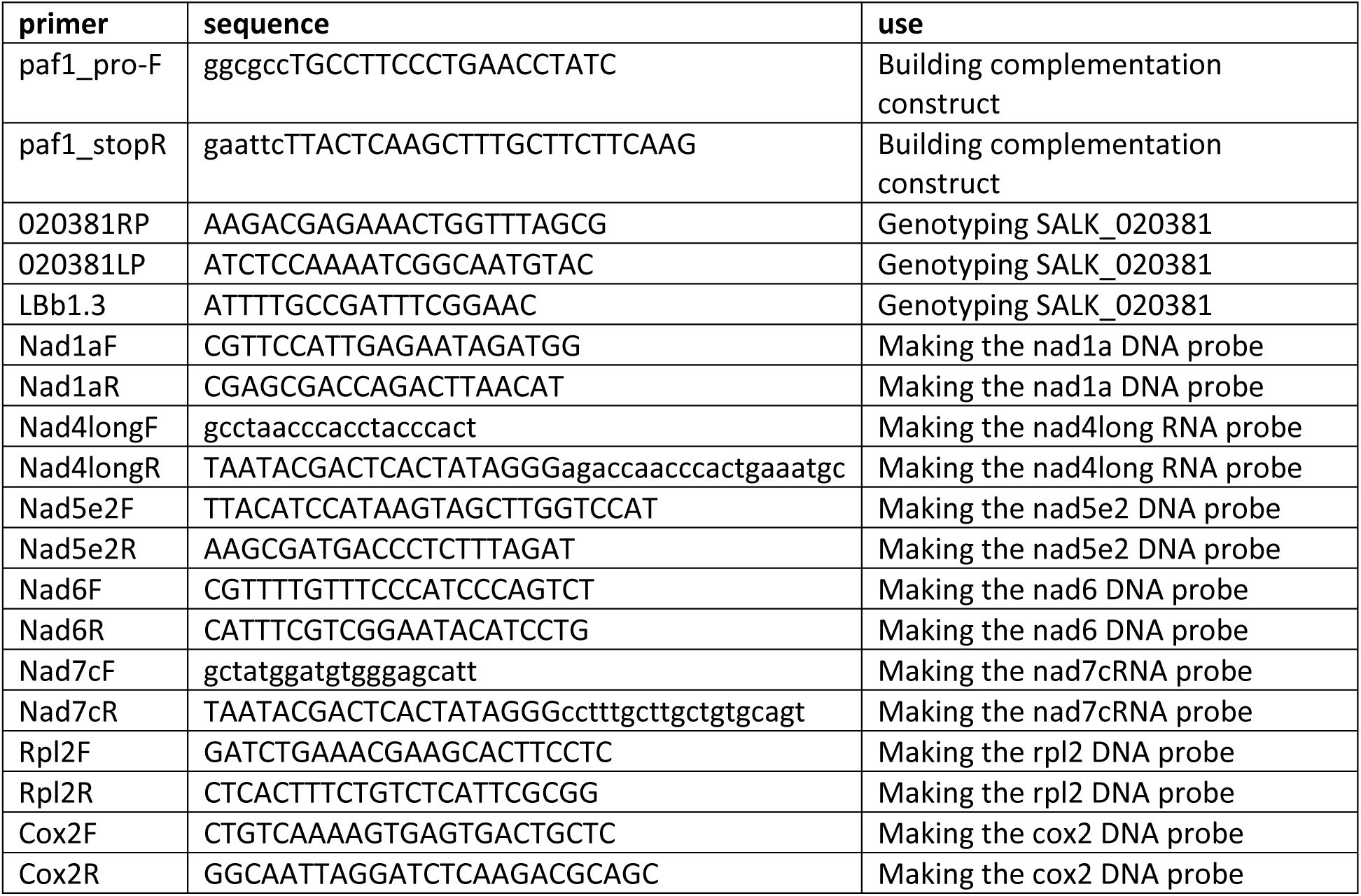
Primers used in this study.

## References

Adams KL, Palmer JD (2003) Evolution of mitochondrial gene content: gene loss and transfer to the nucleus. Mol Phylogenet Evol 29: 380–95

Allen JW (2011) Cytochrome c biogenesis in mitochondria--Systems III and V. FEBS J 278: 4198–216

Boyes DC, Zayed AM, Ascenzi R, McCaskill AJ, Hoffman NE, Davis KR, Gorlach J (2001) Growth stage-based phenotypic analysis of Arabidopsis: a model for high throughput functional genomics in plants. Plant Cell 13: 1499–510

Coutu C, Brandle J, Brown D, Brown K, Miki B, Simmonds J, Hegedus DD (2007) pORE: a modular binary vector series suited for both monocot and dicot plant transformation. Transgenic Res 16: 771–81

Cox J, Mann M (2008) MaxQuant enables high peptide identification rates, individualized p.p.b.-range mass accuracies and proteome-wide protein quantification. Nat Biotechnol 26: 1367–72

Elurbe DM, Huynen MA (2016) The origin of the supernumerary subunits and assembly factors of complex I: A treasure trove of pathway evolution. Biochim Biophys Acta 1857: 971–9

Eme L, Tamarit D, Caceres EF, Stairs CW, De Anda V, Schon ME, Seitz KW, Dombrowski N, Lewis WH, Homa F, Saw JH, Lombard J, Nunoura T, Li WJ, Hua ZS, Chen LX, Banfield JF, John ES, Reysenbach AL, Stott MB et al. (2023) Inference and reconstruction of the heimdallarchaeial ancestry of eukaryotes. Nature 618: 992–999

Eubel H, Braun HP, Millar AH (2005) Blue-native PAGE in plants: a tool in analysis of protein-protein interactions. Plant Methods 1: 11

Formosa LE, Dibley MG, Stroud DA, Ryan MT (2018) Building a complex complex: Assembly of mitochondrial respiratory chain complex I. Semin Cell Dev Biol 76: 154–162

Formosa LE, Mimaki M, Frazier AE, McKenzie M, Stait TL, Thorburn DR, Stroud DA, Ryan MT (2015) Characterization of mitochondrial FOXRED1 in the assembly of respiratory chain complex I. Hum Mol Genet 24: 2952–65

Fuchs P, Rugen N, Carrie C, Elsasser M, Finkemeier I, Giese J, Hildebrandt TM, Kuhn K, Maurino VG, Ruberti C, Schallenberg-Rudinger M, Steinbeck J, Braun HP, Eubel H, Meyer EH, Muller-Schussele SJ, Schwarzlander M (2020) Single organelle function and organization as estimated from Arabidopsis mitochondrial proteomics. Plant J 101: 420–441

Gray MW (2012) Mitochondrial evolution. Cold Spring Harb Perspect Biol 4: a011403

Gualberto JM, Le Ret M, Beator B, Kuhn K (2015) The RAD52-like protein ODB1 is required for the efficient excision of two mitochondrial introns spliced via first-step hydrolysis. Nucleic Acids Res 43: 6500–10

Guerrero-Castillo S, Baertling F, Kownatzki D, Wessels HJ, Arnold S, Brandt U, Nijtmans L (2017) The Assembly Pathway of Mitochondrial Respiratory Chain Complex I. Cell Metab 25: 128–139

Hansen BO, Meyer EH, Ferrari C, Vaid N, Movahedi S, Vandepoele K, Nikoloski Z, Mutwil M (2018) Ensemble gene function prediction database reveals genes important for complex I formation in Arabidopsis thaliana. New Phytol 217: 1521–1534

Hirst J (2013) Mitochondrial complex I. Annu Rev Biochem 82: 551–75

Hooper CM, Castleden IR, Tanz SK, Aryamanesh N, Millar AH (2017) SUBA4: the interactive data analysis centre for Arabidopsis subcellular protein locations. Nucleic Acids Res 45: D1064–D1074

Hunte C, Zickermann V, Brandt U (2010) Functional modules and structural basis of conformational coupling in mitochondrial complex I. Science 329: 448–51

Ivanova A, Gill-Hille M, Huang S, Branca RM, Kmiec B, Teixeira PF, Lehtio J, Whelan J, Murcha MW (2019) A Mitochondrial LYR Protein Is Required for Complex I Assembly. Plant Physiol 181: 1632–1650

Jumper J, Evans R, Pritzel A, Green T, Figurnov M, Ronneberger O, Tunyasuvunakool K, Bates R, Zidek A, Potapenko A, Bridgland A, Meyer C, Kohl SAA, Ballard AJ, Cowie A, Romera-Paredes B, Nikolov S, Jain R, Adler J, Back T et al. (2021) Highly accurate protein structure prediction with AlphaFold. Nature 596: 583–589

Keeling PJ, Eglit Y (2023) Openly available illustrations as tools to describe eukaryotic microbial diversity. PLoS Biol 21: e3002395

Klusch N, Dreimann M, Senkler J, Rugen N, Kuhlbrandt W, Braun HP (2023) Cryo-EM structure of the respiratory I + III(2) supercomplex from Arabidopsis thaliana at 2 A resolution. Nature plants 9: 142–156

Klusch N, Senkler J, Yildiz O, Kuhlbrandt W, Braun HP (2021) A ferredoxin bridge connects the two arms of plant mitochondrial complex I. Plant Cell 33: 2072–2091

Laemmli UK (1970) Cleavage of structural proteins during the assembly of the head of bacteriophage T4. Nature 227: 680–5

Laube E, Schiller J, Zickermann V, Vonck J (2024) Using cryo-EM to understand the assembly pathway of respiratory complex I. Acta Crystallogr D Struct Biol 80: 159–173

Lemire BD (2015) Evolution of FOXRED1, an FAD-dependent oxidoreductase necessary for NADH:ubiquinone oxidoreductase (Complex I) assembly. Biochim Biophys Acta 1847: 451–457

Ligas J, Pineau E, Bock R, Huynen MA, Meyer EH (2019) The assembly pathway of complex I in Arabidopsis thaliana. Plant J 97: 447–459

Maclean AE, Hertle AP, Ligas J, Bock R, Balk J, Meyer EH (2018) Absence of Complex I Is Associated with Diminished Respiratory Chain Function in European Mistletoe. Curr Biol 28: 1614–1619 e3

Maldonado M, Fan Z, Abe KM, Letts JA (2023) Plant-specific features of respiratory supercomplex I + III(2) from Vigna radiata. Nature plants 9: 157–168

Maldonado M, Padavannil A, Zhou L, Guo F, Letts JA (2020) Atomic structure of a mitochondrial complex I intermediate from vascular plants. eLife 9: e56664

McKenzie M, Ryan MT (2010) Assembly factors of human mitochondrial complex I and their defects in disease. IUBMB Life 62: 497–502

Meyer EH, Letts JA, Maldonado M (2022) Structural insights into the assembly and the function of the plant oxidative phosphorylation system. New Phytol 235: 1315–1329

Meyer EH, Tomaz T, Carroll AJ, Estavillo G, Delannoy E, Tanz SK, Small ID, Pogson BJ, Millar AH (2009) Remodeled respiration in ndufs4 with low phosphorylation efficiency suppresses Arabidopsis germination and growth and alters control of metabolism at night. Plant Physiol 151: 603–19

Meyer EH, Welchen E, Carrie C (2019) Assembly of the Complexes of the Oxidative Phosphorylation System in Land Plant Mitochondria. Annu Rev Plant Biol 70: 23–50

Mirdita M, Schutze K, Moriwaki Y, Heo L, Ovchinnikov S, Steinegger M (2022) ColabFold: making protein folding accessible to all. Nat Methods 19: 679–682

Nietzel T, Mostertz J, Ruberti C, Nee G, Fuchs P, Wagner S, Moseler A, Muller-Schussele SJ, Benamar A, Poschet G, Buttner M, Moller IM, Lillig CH, Macherel D, Wirtz M, Hell R, Finkemeier I, Meyer AJ, Hochgrafe F, Schwarzlander M (2020) Redox-mediated kick-start of mitochondrial energy metabolism drives resource-efficient seed germination. Proc Natl Acad Sci U S A 117: 741–751

Padavannil A, Ayala-Hernandez MG, Castellanos-Silva EA, Letts JA (2022) The Mysterious Multitude: Structural Perspective on the Accessory Subunits of Respiratory Complex I. Front Mol Biosci 8

Parey K, Wirth C, Vonck J, Zickermann V (2020) Respiratory complex I - structure, mechanism and evolution. Curr Opin Struct Biol 63: 1–9

Perez-Riverol Y, Bandla C, Kundu DJ, Kamatchinathan S, Bai J, Hewapathirana S, John NS, Prakash A, Walzer M, Wang S, Vizcaino JA (2025) The PRIDE database at 20 years: 2025 update. Nucleic Acids Res 53: D543–D553

Roger AJ, Munoz-Gomez SA, Kamikawa R (2017) The Origin and Diversification of Mitochondria. Curr Biol 27: R1177–R1192

Rohricht H, Przybyla-Toscano J, Forner J, Boussardon C, Keech O, Rouhier N, Meyer EH (2023) Mitochondrial ferredoxin-like is essential for forming complex I-containing supercomplexes in Arabidopsis. Plant Physiol 191: 2170–2184

Rugen N, Schaarschmidt F, Eirich J, Finkemeier I, Braun HP, Eubel H (2021) Protein interaction patterns in Arabidopsis thaliana leaf mitochondria change in dependence to light. Biochim Biophys Acta Bioenerg 1862: 148443

Sanchez-Caballero L, Elurbe DM, Baertling F, Guerrero-Castillo S, van den Brand M, van Strien J, van Dam TJP, Rodenburg R, Brandt U, Huynen MA, Nijtmans LGJ (2020) TMEM70 functions in the assembly of complexes I and V. Biochim Biophys Acta Bioenerg 1861: 148202

Sanchez-Caballero L, Guerrero-Castillo S, Nijtmans L (2016) Unraveling the complexity of mitochondrial complex I assembly: A dynamic process. Biochim Biophys Acta 1857: 980–90

Schäfer K, Kunzler P, Schneider K, Klingl A, Eubel H, Carrie C (2020) The Plant Mitochondrial TAT Pathway Is Essential for Complex III Biogenesis. Curr Biol 30: 840–853 e5

Schertl P, Sunderhaus S, Klodmann J, Grozeff GE, Bartoli CG, Braun HP (2012) L-galactono-1,4-lactone dehydrogenase (GLDH) forms part of three subcomplexes of mitochondrial complex I in Arabidopsis thaliana. J Biol Chem 287: 14412–9

Schimmeyer J, Bock R, Meyer EH (2016) L-Galactono-1,4-lactone dehydrogenase is an assembly factor of the membrane arm of mitochondrial complex I in Arabidopsis. Plant Mol Biol 90: 117–26

Schroder L, Hohnjec N, Senkler M, Senkler J, Kuster H, Braun HP (2022) The gene space of European mistletoe (Viscum album). Plant J 109: 278–294

Senkler J, Rugen N, Eubel H, Hegermann J, Braun HP (2018) Absence of Complex I Implicates Rearrangement of the Respiratory Chain in European Mistletoe. Curr Biol 28: 1606–1613 e4

Senkler J, Senkler M, Eubel H, Hildebrandt T, Lengwenus C, Schertl P, Schwarzlander M, Wagner S, Wittig I, Braun HP (2017) The mitochondrial complexome of Arabidopsis thaliana. Plant J 89: 1079–1092

Signes A, Fernandez-Vizarra E (2018) Assembly of mammalian oxidative phosphorylation complexes I-V and supercomplexes. Essays Biochem 62: 255–270

Smirnoff N, Wheeler GL (2000) Ascorbic acid in plants: biosynthesis and function. Crit Rev Biochem Mol Biol 35: 291–314

Soufari H, Parrot C, Kuhn L, Waltz F, Hashem Y (2020) Specific features and assembly of the plant mitochondrial complex I revealed by cryo-EM. Nat Commun 11: 5195

Stroud DA, Surgenor EE, Formosa LE, Reljic B, Frazier AE, Dibley MG, Osellame LD, Stait T, Beilharz TH, Thorburn DR, Salim A, Ryan MT (2016) Accessory subunits are integral for assembly and function of human mitochondrial complex I. Nature 538: 123–126

Tan YF, Millar AH, Taylor NL (2012) Components of mitochondrial oxidative phosphorylation vary in abundance following exposure to cold and chemical stresses. J Proteome Res 11: 3860–79

Tyanova S, Temu T, Sinitcyn P, Carlson A, Hein MY, Geiger T, Mann M, Cox J (2016) The Perseus computational platform for comprehensive analysis of (prote)omics data. Nat Methods 13: 731–40

Veeraragavan S, Johansen M, Johnston IG (2024) Evolution and maintenance of mtDNA gene content across eukaryotes. Biochem J 481: 1015–1042

Wagener N, Neupert W (2012) Bcs1, a AAA protein of the mitochondria with a role in the biogenesis of the respiratory chain. J Struct Biol 179: 121–5

Wheeler G, Ishikawa T, Pornsaksit V, Smirnoff N (2015) Evolution of alternative biosynthetic pathways for vitamin C following plastid acquisition in photosynthetic eukaryotes. eLife 4

Yip CY, Harbour ME, Jayawardena K, Fearnley IM, Sazanov LA (2011) Evolution of respiratory complex I: “supernumerary” subunits are present in the alpha-proteobacterial enzyme. J Biol Chem 286: 5023–33

